# Sub-second analysis of locomotor activity in Parkinsonian mice

**DOI:** 10.1101/2024.12.26.630411

**Authors:** Daniil Berezhnoi, Hiba Douja Chehade, Gabriel Simms, Liqiang Chen, Kishore Kumar S. Narasimhan, Shashank M. Dravid, Hong-Yuan Chu

## Abstract

The degeneration of midbrain dopamine (DA) neurons disrupts the neural control of natural behavior, such as walking, posture, and gait in Parkinson’s disease. While some aspects of motor symptoms can be managed by dopamine replacement therapies, others respond poorly. Recent advancements in machine learning-based technologies offer opportunities to better understand the organizing principles of behavior modules at fine time scales and its dependence on dopaminergic modulation. In the present study, we applied the motion sequencing (MoSeq) platform to study the spontaneous locomotor activities of neurotoxin and genetic mouse models of Parkinsonism as the midbrain DA neurons progressively degenerate. We also evaluated the treatment efficacy of levodopa (L-DOPA) on behavioral modules at fine time scales. We revealed robust changes in the kinematics and usage of the behavioral modules that encode spontaneous locomotor activity. Further analysis demonstrates that fast behavioral modules with higher velocities were more vulnerable to loss of DA and preferentially affected at early stages of Parkinsonism. Last, L-DOPA effectively improved the velocity, but not the usage and transition probability, of behavioral modules in Parkinsonian animals. In conclusion, the hypokinetic phenotypes in Parkinsonism involve the decreased velocities of behavioral modules and their disrupted temporal organization during movement. Moreover, we showed that the therapeutic effect of L-DOPA is mainly mediated by its effect on the velocities of behavior modules at fine time scales. This work documents robust changes in the velocity, usage, and temporal organization of behavioral modules and their responsiveness to dopaminergic treatment under the Parkinsonian state.

**Significance Statement:** Parkinson’s disease is the second largest neurodegenerative disease without a cure. Detection of subtle Parkinsonian signs is critical for disease-modification by applying early interventions. The present work explores the possibility of using machine learning-based approaches for early detection of subtle behavioral changes in Parkinsonian animals and evaluating the therapeutic efficacy of dopaminergic medications.

## Introduction

Dr. James Parkinson noted Parkinsonian signs by observing individuals on the street and documented the impaired locomotor activities and gait control in patients’ daily living ^1^. Over decades, the diagnosis of Parkinson’s disease (PD) has been primarily conducted in clinical settings, where trained clinicians assess gross motor and cognitive function as well as mental status ^2^. However, when and how PD alters patients’ behavior in their natural environment remains poorly understood. This is a critical question to address, considering that PD is a chronic and progressive neurological disorder. Detection of motor and nonmotor signs associated with early PD can provide valuable opportunities for effective interventions to alleviate clinical symptoms and even slow down the disease progression. It is established that degeneration of dopaminergic neurons in the substantia nigra pars compacta (SNc) results in hypokinetic symptoms, including akinesia and bradykinesia. Accordingly, restoration of dopamine (DA) levels in the brain by levodopa (L-DOPA) administration can effectively manage hypokinetic symptoms in patients ^1^. With emerging machine-learning-based technologies, it becomes possible to use unbiased and data-driven approaches for detailed analysis of motor deficits and the efficacy of dopaminergic medications in PD at fine time scales ^3,4^. Such information could reveal new behavioral biomarkers for unbiased PD diagnosis and facilitate the design of effective treatment.

Naturalistic motor behavior is generated by orchestrating distinct behavioral modules with different kinematic properties and probability of expression into meaningful actions ^5^. Recent reports demonstrated that DA levels in the striatum and activities of striatal projection neurons play a critical role in representing and orchestrating behavioral modules into action sequences ^6,7^. Building on these observations, decreased striatal DA levels in PD are predicted to alter the kinematics, the probability of expression, and the organization of behavioral modules into actions, leading to the progressive manifestation of hypokinetic symptoms. In addition, while L-DOPA can effectively improve the motor activity of PD patients, whether L-DOPA restores the basic principles of orchestrating behavioral modules into naturalistic movements remains to be determined.

In the present study, we hypothesized that loss of DA decreases the kinematic measures and frequencies of usage of behavioral modules of naturalistic locomotor activity and that the magnitude of such reduction scales up as the level of striatal DA dampens in PD. We further hypothesized that L-DOPA can restore kinematics and usage of behavioral modules, thereby contributing to its beneficial effects in PD. Here, we employed machine learning-based technologies to perform unsupervised and unbiased segmentation and quantification of spontaneous behavior of two established mouse models of Parkinsonism to test these hypotheses.

## Material and Methods

### Animals

Adult C57BL/6 mice (JAX#: 000664, The Jackson Laboratory; RRID: IMSR_JAX:000664) of both sexes at 3-5 months were used for 6-hydroxydopamine (6-OHDA) lesion studies. Mitochondrial transcription factor A (*Tfam*) floxed mice (*Tfam*^loxP/loxP^, JAX#: 026123, The Jackson Laboratory; RRID: IMSR_JAX:026123) and *Dat*-Cre (JAX#: 006660, The Jackson Laboratory; RRID: IMSR_JAX:006660) mice were used to generate the mitoPark mice, as a model of progressive nigrostriatal degeneration ^8–10^. Specifically, the double heterozygous mice (*Dat-Cre*^Cre/+^ x *Tfam*^loxP/+^) were crossed with *Tfam*^loxP/loxP^ mice to selectively delete *Tfam* in dopamine neurons, resulting in adult-onset progressive nigrostriatal degeneration and parkinsonian motor deficits. Age-matched littermates with *Tfam^l^*^oxP/loxP^ or *Tfam*^loxP/+^ genotypes were used as controls. Mice were housed up to four animals per cage under a 12/12 h light/dark cycle with access to food and water *ad libitum* in accordance with NIH guidelines for the care and use of animals. All the animal experiments were reviewed and approved by IACUC at Van Andel Institute (AUP# 22-02-006).

### Stereotaxic surgery

To induce degeneration of nigral DA neurons, we injected 6-OHDA (3–4 mg/ml, 1.0 μl) unilaterally into the medial forebrain bundle (MFB; anteroposterior = −0.7 mm; mediolateral = +1.2 mm; dorsoventral = −4.7 mm) under 2% isoflurane anesthesia over 10 min using a 10-μl syringe (Hamilton). Control animals were injected with HEPES-buffered saline (HBS) in the same location. Desipramine (25 mg/kg) and pargyline (50 mg/kg) were subcutaneously injected 30–40 min before 6-OHDA injection to reduce damage to the noradrenergic system and enhance the toxicity of 6-OHDA. Mice were allowed to recover after surgery in a heated cage with access to food and water and closely monitored until the behavioral studies.

### Behavioral experiments

All behavioral experiments were conducted 3-5 weeks post 6-OHDA or HBS injections. To study progressive changes in the spontaneous behavior during gradual loss of striatal DA, we recorded the open field locomotor behavior of the mitoPark mice 3 ages representing distinct stages of parkinsonism: at 8 weeks with normal striatal DA levels, at 14 weeks with 50% reduction of the striatal DA levels and early motor signs, and at 24 weeks with a nearly complete loss of striatal DA and severe motor deficits ^10,11^. A separate cohort of mitoPark mice and littermate controls at >24 weeks received L-DOPA (10 mg/kg, i.p.) and benserazide (15 mg/kg, i.p.) injections to evaluate the treatment efficacy of L-DOPA at sub-second levels. All experiments were performed in a room with dimmed lights. Animals were acclimated to the room and the experimenters prior to recordings. In all experiments, we recorded spontaneous animal behavior for 15-20 min in a square (40×40cm) black-colored box with non-reflective walls and a camera mounted on top (DOI: dx.doi.org/10.17504/protocols.io.e6nvwjxmdlmk/v1). A Microsoft Kinect 2.0 camera was used (30 fps) for recordings from 6-OHDA mice and mitoPark mice, while a FLIR Blackfly S camera (Teledyne Technologies, USA) was used for the mitoPark mice plus L-DOPA experiment. See Key Resource Table for details.

### Computational methods

For video analysis, we employed the Keypoint-MoSeq tracking algorithms (V. 0.6.2, https://github.com/dattalab/keypoint-moseq, RRID:SCR_025032) ^12^. Specifically, 1) Keypoint tracking with ResNet50 network pre-trained on a subset of videos using DeepLabCut pipeline (V.2.3.10, https://github.com/DeepLabCut/DeepLabCut, RRID:SCR_021391), 2) egocentric alignment and cropping, leaving only the postural data, 3) principal component analysis (PCA) to reduce the dimensionality of the resulting time series (5 components for our data), 3) probabilistic modeling fitting autoregressive hidden Markov model (AR-HMM) to the PCA trajectories, 4) fitting full Keypoint-MoSeq model initialized from the AR-HMM, 5) automatic behavioral segmentation and analysis using the motion syllables assigned by the trained model. All the analyses were performed using the provided pipeline in Jupyter Notebooks (V. 7.1.3, https://jupyter.org, RRID:SCR_018315). Training of the model is done through the process of iterative Gibbs sampling while adjusting the Kappa hyperparameter of the model and leaving all other parameters as defaults (Alpha = 5.7, Gamma = 1e3, Max number of states = 1e2) ^12^.

The motion sequencing platform performs probabilistic analysis of the behavior data, and the results depend on the chosen computational model. We fitted our data to multiple models and compared the results before choosing the final model to be applied to all datasets. To select the model, we performed the kappa scanning procedure on our data, training the models with multiple different kappa values and choosing the one that matched our criteria. We used two main criteria to choose a model for further analysis of behavioral videos. The model should (1) match the target duration of the syllable 400 ms (∼12 frames at 30 fps) as described previously ^5^, (2) generate behaviorally meaningful syllables (i.e., consistent behavior modules that an experienced observer can recognize) on the representative grid videos. After the visual screening, the models with inconsistent syllables (e.g., different behavioral modules fall in the same syllable) or too short/long syllables were discarded. Velocity of syllables is calculated by the MoSeq algorithm using the 2d coordinates (x-y) coordinates of the single point (i.e., animal centroid) without including z-axis information. At the same time, MoSeq algorithm assigns a syllable number to each frame. Thus, averaging the differential of the x-y position between all frames assigned to a certain syllable gives the average velocity for that syllable. For conventional locomotor measurement, the velocity is calculated from the x-y coordinates in the MoSeq data frame as the average per the session by dividing the total distance traveled to the time of the session in seconds. For further reference on the exact codes, see the documentation available online (https://keypoint-moseq.readthedocs.io/en/stable/index.html).

### Immunohistochemical assessment of striatal DA depletion

We used western blot and immunofluorescence staining against tyrosine hydroxylase (TH) in the striatum to assess the level of DA depletion in 6-OHDA mice and the 24 weeks old mitoPark mice, respectively, at the end of this study. 70 μm thick brain sections were prepared using a vibratome (VT1000S; Leica microsystems Inc.). Immunochemical detection of TH was conducted in PBS containing 0.2% Triton X-100 (Fisher Scientific) and 2% normal donkey serum (Sigma-Aldrich). Brain sections were incubated in the primary antibody (mouse anti-TH antibody, 1:2000, catalog #MAB318, Sigma-Aldrich, RRID:AB_2313764) for 48 h at 4°C or overnight at room temperature, washed in PBS, and then incubated in the secondary antibody (donkey anti-mouse Alexa Fluor 594, catalog #715-585-150, RRID:AB_2340846) for 90 min at room temperature before washing with PBS. Brain sections were mounted on glass slides using VECTASHIELD antifade mounting medium (catalog #H-1000, Vector Laboratories) and were cover-slipped. TH immunoreactivity in the dorsal striatum and the overlying motor cortex in the same section was imaged using an Olympus BX63F microscope equipped with an Olympus DP80 camera or a confocal laser scanning microscope (A1R; Nikon). The intensity of TH immunofluorescence was quantified using ImageJ (NIH). The ratio of TH-immunoreactivity (TH-ir) between the dorsal striatum and the motor cortex was determined to assess the striatal DA depletion in mitoPark mice.

<u>Tissue lysate preparation and Immunoblotting.</u> 6-OHDA mice were anesthetized with isoflurane and then decapitated. The brain was removed from the skull and rinsed with ice-cold ACSF to remove any adherent blood. One millimeter (1 mm) coronal sections were taken using brain matrix and the dorsal striatum (dSTR) was punched from the lesioned hemisphere. We used the Allen Brain Atlas: Mouse brain (https://mouse.brain-map.org/static/atlas) as an anatomical reference atlas for punching out the dSTR. The punched tissue was processed for both whole tissue lysate using RIPA buffer (total protein) as well as synaptoneurosomes (fraction containing enriched synaptic components) as previously described ^13^. For synaptoneurosomes preparation, dSTR was homogenized in a synaptoneurosome buffer (10 mM HEPES, 1 mM EDTA, 2 mM EGTA, 0.5 mM DTT, 0.5 mM PMSF, 50 μg/ml soy-bean trypsin inhibitor, 0.25% of phosphatase inhibitor cocktail 2% and 3%, and 0.25% of protease inhibitor cocktail) and sonicated with 3 pulses (FB50, Fisher Scientific, Pittsburgh, PA, USA). The resulting sample was filtered twice through three layers of a pre-wetted 100 μm pore nylon filter (CMN-0105-D, Small Parts Inc., Logansport, IN, USA) held in a 13 mm diameter filter holder (XX3001200, Milipore, MA, USA). Then the resulting filtrate was filtered once through a pre-wetted 5 μm pore hydrophilic filter (CMN-0005-D, Small Parts Inc.). The filtrate was centrifuged at 1000 × g for 10 min at 4 °C to obtain the synaptoneurosomes fraction. Isolated synaptoneurosomes were resuspended in synaptoneurosomes buffer containing 0.32 M sucrose, and 1 mM NaHCO3, stored in –80°C until further use. Protein concentration of RIPA lysates and synaptoneurosomes was measured with Pierce™ BCA Protein Assay Kit (Cat# 23227, Thermo Fisher Scientific).

Immunoblotting was performed as previously described^13^. Briefly, RIPA lysates and synaptoneurosomes were loaded on 10% sodium dodecyl sulfate gel in equal amounts (50 μg/well). The samples were run at 100 volts for 90 min. Gels were transferred to activated (incubation in methanol) polyvinylidene fluoride (PVDF) membrane (GE Healthcare, Piscataway, NJ, USA) at 115 volts for 1 h 45 min. Transfer was blocked with 5% milk in Tris-buffered Saline with 0.1% Tween 20 (TBST) for 1 h at room temperature. The membranes were kept overnight for incubation in primary antibodies including mouse monoclonal anti-tyrosine hydroxylase (1:2000, BioLegend Cat# 818002, RRID: AB_2734573) and mouse monoclonal anti-GAPDH (1:4000, EMD Millipore Cat# MAB374, RRID: AB_2107445) at 4°C followed by washing with TBST. Blots were incubated with Peroxidase-AffiniPure Goat Anti-Mouse IgG secondary antibody (Jackson ImmunoResearch Labs Cat# 115-035-003, RRID: AB_10015289) for 1 h at room temperature followed by washing with TBST. Blots were incubated with Super-Signal™ West Pico PLUS chemiluminescent substrate (Cat# 34580, ThermoFisher Scientific, Waltham, MA, USA) and developed in a ChemiDoc imaging system (Bio-Rad Laboratories, Inc., USA). The optical density of each sample was analyzed using ImageJ and normalized to GAPDH. Each western blot data point in each group was obtained from a separate animal.

### Statistical analysis

Data analysis and statistics were performed using Python (Python Programming Language, V.3.9.19, https://www.python.org/downloads/release/python-360, RRID:SCR_008394) and Prism 10 (GraphPad Software, http://www.graphpad.com/,_RRID:SCR_002798)). The main parameters of behavioral syllables (e.g., usage and velocity) were calculated for each animal and then averaged within experimental groups. Probability matrices representing the transitions between syllables were generated for each animal and also averaged within groups to represent the differences in behavioral structure. Entropy values were calculated from transition probability matrices as a sum of local entropies per syllable, as reported by ^5^. Specifically, Let *A* denote the estimated transition matrix, so that *A_ij_* = P[*X*_t+1_ = *j* | *X*_t_ = i] for *i,j* = 1,2,…, *n* and all time indices *t*, and let π denote the steady-state distribution of the Markov chain, so that π_i_ = Pr[*X*_t_ = i]. The entropy rate on the transition matrix was calculated as ∑i,j π_i_ *A_ij_ ln A_ij_*. Data are reported as the mean and standard error of the mean (SEM). Kruskal-Wallis test with post-hoc Dunn’s test in the MoSeq package was used to compare the velocity and usage of behavioral syllables across conditions. MoSeq packages corrected raw P values for multiple comparisons using the Benjamini-Hochberg methods, with a false discovery rate (FDR) of 0.05 or less. Mann-Whitney U test was used to compare the statistical significance of the conventional measure of locomotor activity (i.e., velocity) and striatal TH levels between groups, and α < 0.05 was considered statistically significant. Details of behavioral analysis can be found at: https://github.com/ChuLab-GU/behavior-analysis

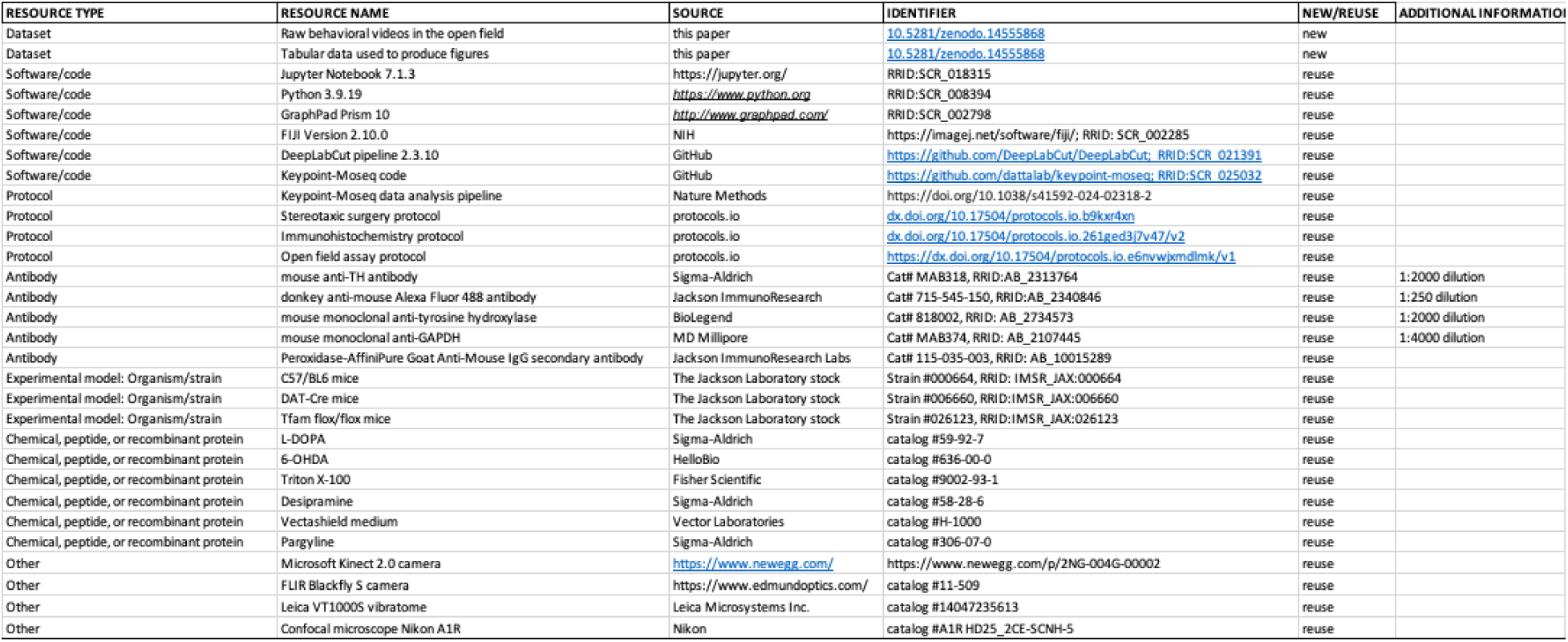
Key Resource Table.

## Results

### Loss of DA alters the velocity and usage of behavioral modules

First, we employed mice with 6-hydroxydopamine (6-OHDA) lesions as a model of nigrostriatal degeneration, which is known to recapitulate behavioral phenotypes and pathophysiological features of human PD and has been widely used for preclinical PD research ^14–19^. To model an advanced Parkinsonian state with nearly complete DA depletion, we injected 6-OHDA into the MFB to induce degeneration of the nigrostriatal pathway and nearly complete striatal DA depletion (**Figure 1A, B**) ^15,17,20^. Three weeks post-surgery, locomotor activities of 6-OHDA– and vehicle-injected mice (i.e., 6-OHDA mice and controls, respectively) were recorded for 20 min in an open field arena. Analysis of conventional locomotor measures demonstrates that 6-OHDA mice showed a significant reduction of general motor activity relative to controls (e.g., velocity, control = 8.9±0.6 cm/sec, n = 6 mice; 6-OHDA = 5.5±0.2 cm/sec, n = 7 mice; P < 0.001). These results confirm the severe Parkinsonian motor deficits following the degeneration of the dopaminergic nigrostriatal pathway.

**Figure 1.**
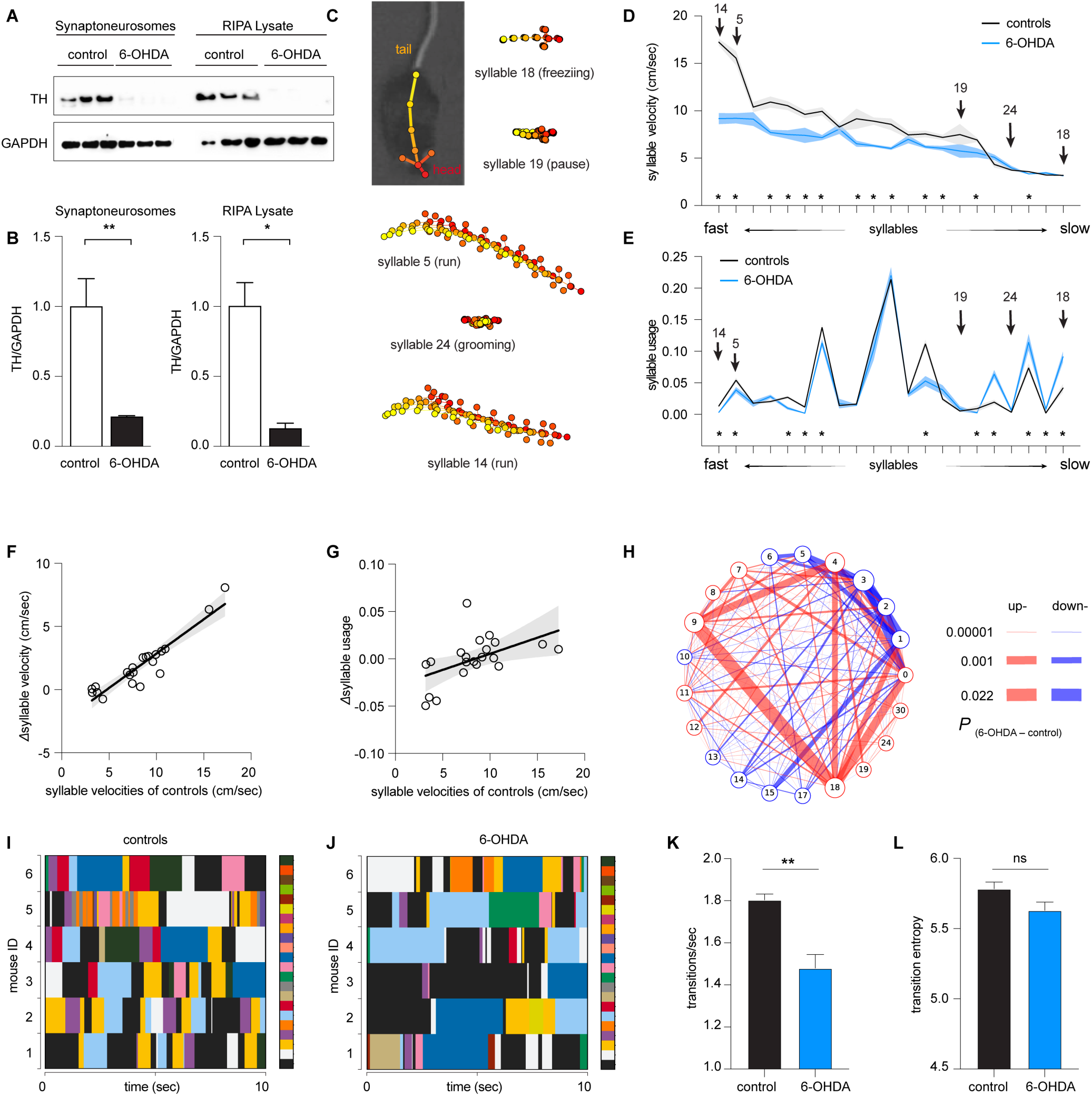
Loss of DA alters the velocity and usage of behavioral modules of 6-OHDA mice. **A**) Western blotting analysis of synaptoneurosome and RIPA lysate TH protein from the dorsal striatum of control and 6-OHDA mice. A significant reduction of striatal TH protein as seen in both synaptoneurosome (p = 0.0023, n = 3 mice/group, Welch’s t test) and RIPA lysate (p = 0.03, n = 3mice/group, Welch’s t test). **C**) Representative behavioral syllables identified by MoSeq. **D-E**) Syllable velocity (D) and usage (E) of 6-OHDA mice and controls ranked by velocities for visualization. Arrows indicated the representative syllables in (C). Asterisks indicate syllables showing statistical significance in either the syllable velocities (D) or usage (E) between 6-OHDA mice and controls. See Table 1 for the source data. **F-G**) Differences in syllable velocities (Δvelocity, D) or usage (Δusage, E) between controls and 6-OHDA mice were plotted against the syllable velocities in controls. **H**) Transition graphs representing the changes in behavioral structure as an increase (red) or decrease (blue) in the probability of transition between numbered syllables in 6-OHDA mice relative to controls. I-J) Representative plots showing the temporal organization of spontaneous behavior in controls (I) and 6-OHDA mice (J). Different colors represent distinct behavioral syllables. Only six 6-OHDA mice were shown in the representative plot. **K**) Summarized bar plot showing the reduced transition frequency of syllables in 6-OHDA mice relative to controls (control = 1.8±0.03, n = 6 mice; 6-OHDA = 1.48±0.07, n = 7 mice, P = 0.008, Mann-Whitney U test). L) Summarized bar plot showing a nonsignificant change in the transition entropy of syllables in 6-OHDA mice relative to controls (control = 5.78±0.04, n = 6 mice; 6-OHDA = 5.63±0.06, n = 7 mice, P = 0.1, Mann-Whitney U test).

Next, we employed the motion sequencing (i.e., MoSeq) approach to interrogate the locomotor activity of Parkinsonian mice at fine timescales. MoSeq automatically identifies behavioral modules at sub-second levels (i.e., syllables) and determines the probability of their usage in locomotor activities ^3–7,12,21^. MoSeq recognized morphologically distinct sub-second behavioral syllables with comparable mean durations of 588±5.7 ms for controls and 538±6.3 for 6-OHDA mice. Αmong all the identified behavioral modules, 21 syllables were expressed more than 0.5% of the time (see examples in **Figure 1C**). Each syllable could be described as a behavior module by a human observer, including various pauses (e.g., syllable 19), freezing (i.e., no change in body location for 1 sec or more, syllable 18), run (e.g., syllable 5 and 14), grooming (e.g., syllable 24), rearing, and turns. Although unilateral turnings are behavioral features of mice with 6-OHDA lesion, they were not identified as a single syllable by Moseq. This might be because spontaneous unilateral turnings in an open field arena usually take longer than a second. In the rest of the study, we focused on the 21 most frequently used and meaningful behavior modules to characterize Parkinsonian phenotypes in mice at the sub-second level.

For the purpose of visualization, we ranked syllables based on their velocities, ranging from faster run-like syllables to slower freezing-like syllables (**Figure 1C-D**). We found a significant decrease in the velocities of faster syllables of 6-OHDA mice relative to controls (e.g., the velocity of syllable# 14 [i.e., run], controls = 17.2±0.36 cm/sec, n = 6 mice; 6-OHDA = 9.2±0.6 cm/sec, n = 7 mice, P < 0.0001, **Figure 1D**). In contrast, the velocities of most slower syllables (e.g., pause and freezing) were not altered between groups (e.g., the velocities of syllable#18 [i.e., freezing], controls = 3.19±0.045 cm/sec, 6-OHDA mice = 3.14±0.019 cm/sec; the velocities of syllable#19 [i.e., pause], controls = 7.5±1.1 cm/sec, n =6 mice; 6-OHDA mice = 5.76±0.72 cm/sec, n = 7 mice, P > 0.05, **Figure 1D**). Next, we analyzed the frequency of expression (i.e., the usage) of behavioral syllables in 6-OHDA mice and controls. There were significant decreases in the usage of faster syllables and increases in the usage of slower syllables in the 6-OHDA mice relative to controls (**Figure 1E**). For example, 6-OHDA mice used the faster syllable#14 significantly less than controls (controls = 0.013±0.002, n = 6 mice; 6-OHDA = 0.003±0.0008, n = 7 mice; P < 0.0001). In contrast, 6-OHDA mice expressed the slower syllable#18 more frequently than controls (controls = 0.04±0.0065, n = 6 mice; 6-OHDA = 0.09±0.008, n = 7 mice, P = 0.002).

Further, we tested the hypothesis that loss of DA has stronger effects on faster syllables. To do this, we assessed the correlation between the group differences in the velocities and usages [i.e., Δvelocities_(control-6-OHDA)_ and Δusage_(control-6-OHDA)_, respectively) against the syllable velocities of controls (**Figure 1F, G**). Linear regression analysis showed a strong correlation between the Δvelocities and syllable velocities in controls (R^2^ = 0.87, P < 0.0001, **Figure 1F**). Similarly, a statistically significant linear relationship was detected between Δusage and syllable velocities in controls (R^2^ = 0.27, P = 0. 0165, **Figure 1G**).

Together, these results suggest that while the kinematics and usage of behavior modules are altered following the loss of DA, the changes neither uniform nor unidirectional. Specifically, the nigrostriatal dopaminergic degeneration preferentially decreases the kinematics of faster behavioral modules and disrupts the usage of various behavioral modules, resulting in Parkinsonian motor deficits.

### Loss of DA alters the transition probability of behavioral modules

Disrupted usage of behavioral modules is expected to alter the temporal organization of syllables into actions. Indeed, the transition matrix showed that the transition probabilities were up– and down-regulated among subsets behavioral syllables (**Figure 1H**). Moreover, we plotted the temporal organization of behavioral syllables in controls and 6-OHDA mice to further assess the patterns of behavioral module expression and transition. Relative to controls, 6-OHDA mice exhibited prolonged expression of a subgroup of syllables, leading to decreases in syllable transitions (**Figure 1I, J**). We employed entropy (i.e., a measure of disorder level) and the frequency of transitions to quantify the temporal organization of behavioral syllables (**Figure 1K, L**) ^5^. Quantitative analysis confirmed less frequent syllable transitions in 6-OHDA mice than in controls. Specifically, the number of syllable transitions decreased significantly in 6-OHDA mice relative to controls (**Figure 1K**). However, the difference in entropy did not reach statistical significance between 6-OHDA mice and controls (**Figure 1L**). These data indicate that Parkinsonian animals exhibited fewer behavior module transitions and distinct pattern of organizing rules of syllables during naturalistic behavior. This is consistent with a recent study reporting that elevated DA levels in the striatum increase the variabilities of behavioral modules, making them less predictable during naturalistic motor activities ^7^.

Taken together, the above data suggest that hypokinetic symptoms in the Parkinsonian state involve the abnormal sub-second architecture of locomotor behavior, as reflected by 1) the reduced velocities of subsets of syllables, (2) disrupted usage and altered temporal organization of behavioral syllables, and 3) decreased frequency of syllable transitions during spontaneous locomotion activities.

### Time course of velocity changes of behavioral modules in progressive Parkinsonism

PD is a chronic and progressive neurodegenerative disease that affects various aspects of motor control and body movements. Next, we depicted how the observed changes in behavior modules emerge from the early to late stages of Parkinsonism. To address this question, we utilized the mitoPark mice as a model of progressive degeneration of the nigrostriatal dopaminergic pathway (**Figure 2A**) ^8–10,22^. Based on the evidence in the literature, we identified the following age groups of mitoPark mice to study: baseline state at 8 weeks with normal motor function, early motor stage at 14 weeks, and advanced motor stage at 24 weeks ^11^. Spontaneous open-field locomotor activities of mitoPark mice and littermate controls were recorded at 8, 14, and 24 weeks of age. Analysis of locomotory activity using conventional measures confirmed that mitoPark mice showed normal general motor activity at 8 weeks, but exhibited gradual decreases in movement speed from 14 to 24 weeks of age (**Figure 2B**). This time course of Parkinsonism development in mitoPark mice is consistent with previous reports^8–10,22^.

**Figure 2.**
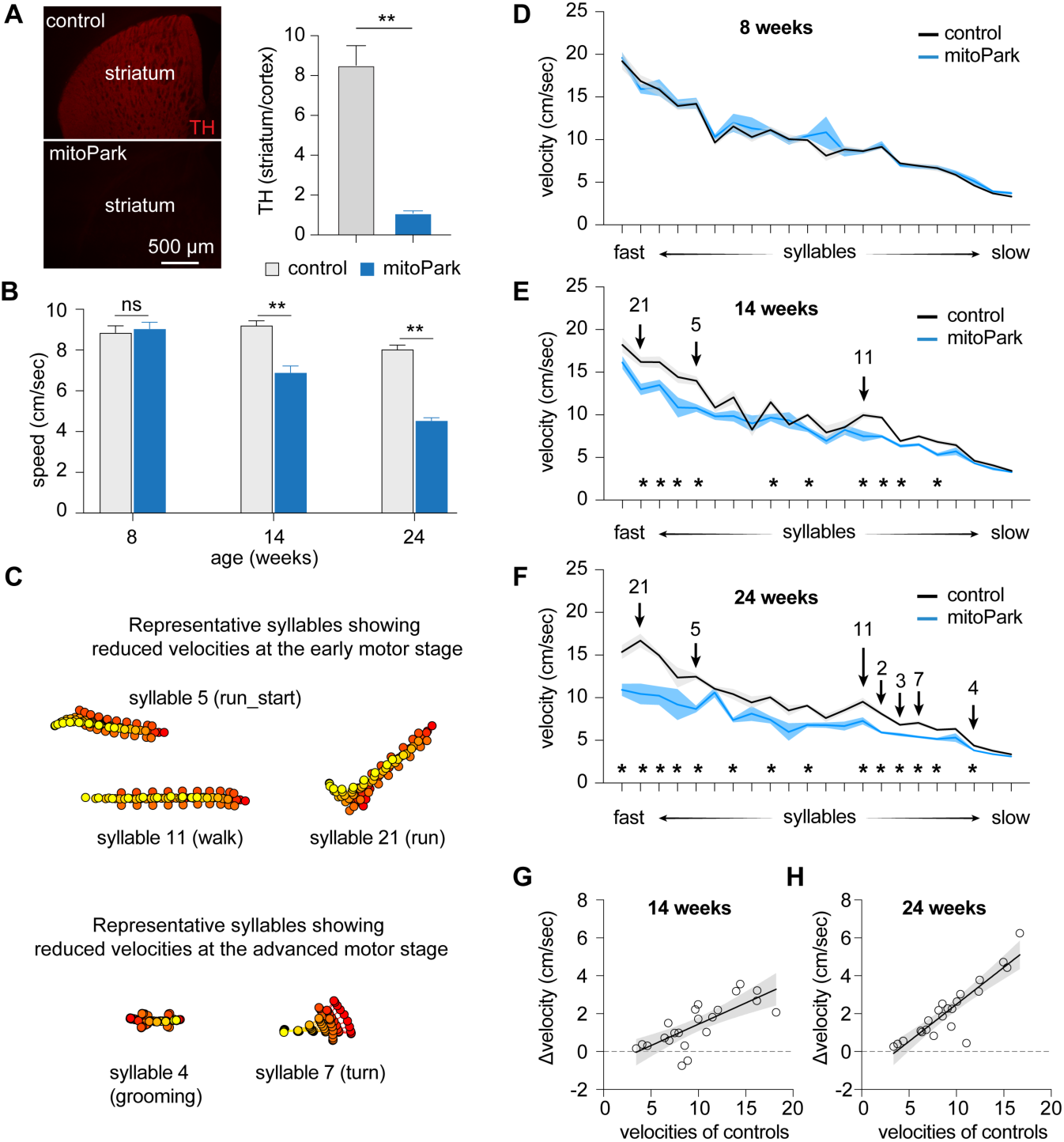
Time course of velocity changes of behavioral modules in mitoPark mice. **A**) mitoPark mice showed a complete loss of striatal TH immunoreactivity at 24 weeks (control = 8.5±0.98, n = 8; mitoPark = 1.1±0.12, n = 7, p < 0.0001). **B**) mitoPark mice exhibited a progressive reduction in general locomotory activity relative to littermate controls Locomotion speed: At 8 weeks, controls = 8.85±0.34 cm/sec, n = 11 mice, mitoPark = 9.04±0.33 cm/sec, n = 11 mice, P = 0.3; At 14 weeks, controls = 9.21±0.23 cm/sec, n = 8 mice, mitoPark = 6.88±0.34 cm/sec, n = 8 mice, P = 0.0002; At 24 weeks, controls = 8.03±0.22 cm/sec, n = 7 mice, mitoPark = 4.53±0.15 cm/sec, n = 9 mice, P = 0.0002. **C**) Representative syllables showing significant changes in syllable velocities at early and late motor stages in mitoPark mice relative to controls. **D-F**) Plots showing progressive reduction of syllable velocities in mitoPark mice relative to age-matched littermate controls at 8-, 14-, and 24-weeks. Arrows indicate the representative syllables shown in (B). Asterisks showing the syllables with statistically significant changes between mitoPark mice and littermate controls. See Table 2 for details. **G-H**) Differences in syllable velocities (Δvelocity) between mitoPark mice and littermate controls were plotted against the syllable velocities in controls.

MoSeq was used to analyze the locomotor activities of mitoPark mice to reveal potential changes in kinematics, usage, and organization of behavior syllables at different age groups. MoSeq identified 99 behavioral syllables expressed by mitoPark mice and littermate controls with a mean duration of 603.5±6.1 ms. 22 out of 99 syllables were used more than 0.5% of the time by animals of both genotypes. Thus, we focused on these 22 most enriched behavioral syllables to characterize the evolvement of the usage and sequence of behavior modules as the nigrostriatal dopaminergic pathway degenerates progressively (**Figure 2C**).

Consistent with the conventional analysis of locomotor activities (**Figure 2B**), mitoPark mice showed no difference in the velocity of various syllables relative to littermate controls at 8 weeks of age (**Figure 2D**). At the early motor stage (i.e., 14 weeks), 10 out of 22 syllables showed reduced velocities in mitoPark mice relative to littermate controls (**Figure 2C, E**). At the advanced motor stage (i.e., 24 weeks), 14 out of 22 syllables showed decreased velocities in mitoPark mice relative to littermate controls (**Figure 2C, F**). Moreover, the syllables with higher velocities (e.g., run-like modules, **Figure 2C**) showed earlier decreases in the velocity, and the magnitude of such changes scaled up from the early to the advanced motor stages. For example, the velocity of the run-like syllable 21 showed 20% reduction at 14 weeks (controls = 16.2±0.59 cm/sec and mitoPark = 12.98±0.67 cm/sec, n = 8 mice/group, P = 0.014); and 38% reduction at 24 weeks (controls = 16.7±0.8 cm/sec, n = 7 mice; mitoPark = 10.45±1.21 cm/sec, n = 8 mice; P = 0.001). As the nigrostriatal dopaminergic pathway degenerated, additional syllables significantly reduced their velocities from the early motor stage to the advanced stages (**Figure 2C, E, F**). Most syllables showing significant changes in velocities at 24 weeks were those involving whole body movements (**Figure 2C, F**), including various turns (e.g., syllable 7), grooming (e.g., syllable 4),

Similar to the mice with 6-OHDA lesion, simple linear regression analysis showed a strong correlation between the Δvelocity (i.e., controls minus mitoPark mice) and syllable velocities of respective controls at 14 and 24 weeks (regression coefficient, 14 weeks = 0.22, P < 0.0001; 24 weeks = 0.39, p < 0.0001) (**Figure 2G-H**). Thus, hypokinetic parkinsonian motor deficits could be partially explained by the progressive reduction of the kinematics of a subset of faster behavioral modules as the nigral DA neurons progressively degenerate.

### Time course of usage changes of behavioral modules in progressive Parkinsonism

When the syllable usage was compared between groups, mitoPark mice gradually showed a subset of differently expressed behavioral modules relative to controls (**Figure 3A-D**). Specifically, the usage of behavioral modules was comparable between controls and mitoPark mice at 8 weeks, except for a transient and slight increase in the usage of syllable 16 by mitoPark mice (**Figure 3B**). At 14 weeks, mitoPark mice showed three additional differently expressed behavioral modules relative to littermate controls, including an upregulation of a freezing-like module (i.e., syllable 9) and a downregulation of a run-like module (e.g., syllable 1). At 24 weeks, mitoPark mice showed 13 differently expressed behavioral modules, which further diverged from the pattern of syllable usage in controls (**Figure 3D**). These changes in syllable usage were not unidirectional, including upregulation of pause-like behavior (e.g., syllable 18) as well as downregulations of faster modules (e.g., syllable 14: run) and the syllables involving large-amplitude whole body movements (e.g., syllable 6: turns). These data suggest that the gradual loss of DA altered the usage of syllables multi-directionally, not uniformly, depending on their distinct roles in organizing naturalistic behavior.

**Figure 3.**
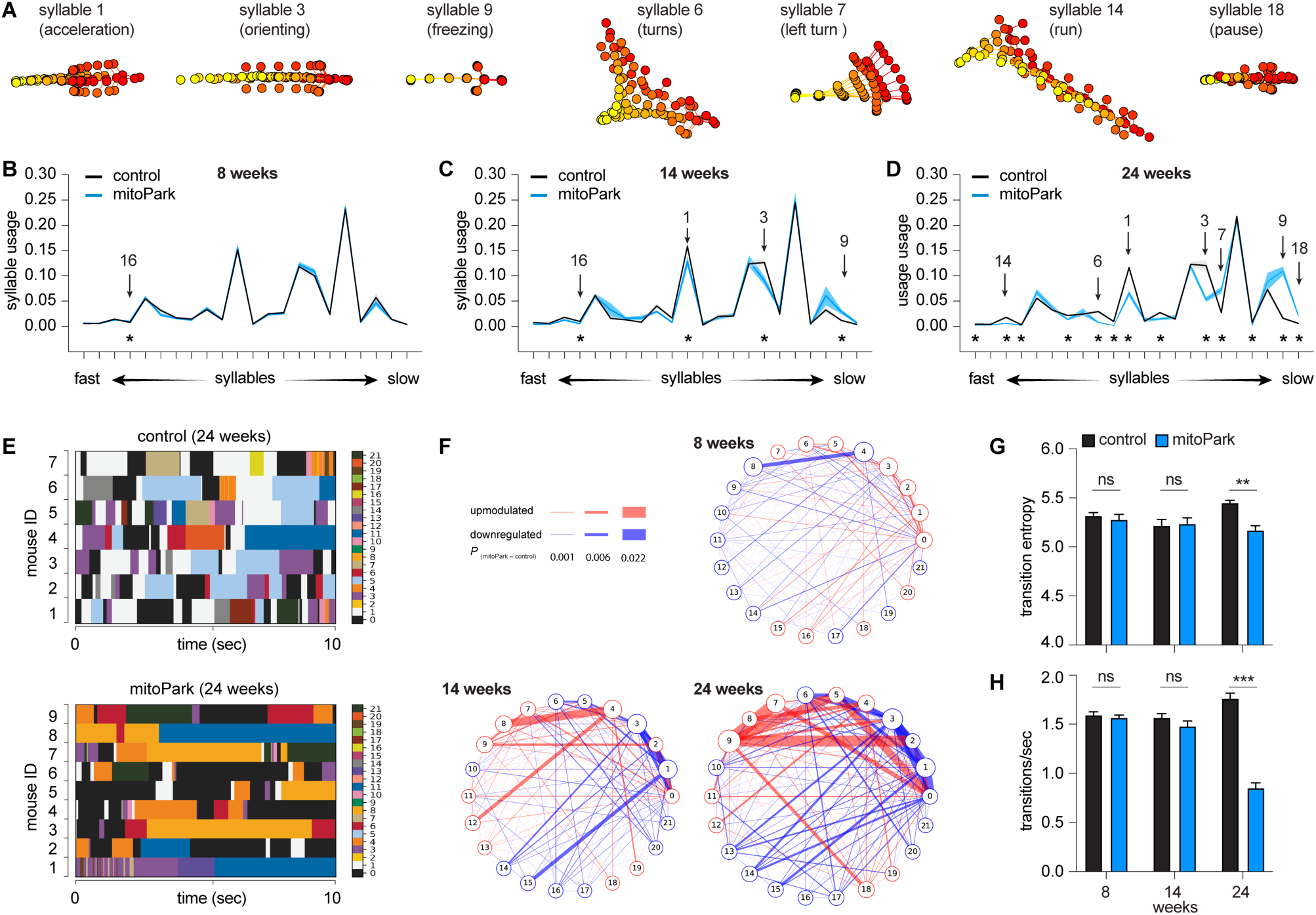
Time course of usage changes of behavioral modules in progressive Parkinsonism. **A**) Representative syllables showing changes in usage in mitoPark relative to littermate controls. **B-D**) Plots of syllable usage between mitoPark mice and littermate controls at 8-, 14-, and 24 weeks. Arrows indicate the representative syllables shown in (A). Asterisks indicate syllables showing statistically significant changes in the usage across time. **E**) Representative plots showing the temporal organization of spontaneous behavior in controls (upper) and mitoPark mice (bottom) at 24 weeks. **F**) Transition graphs representing the changes in behavioral structure as an increase (red) or decrease (blue) in the probability of transition between numbered syllables in mitoPark mice relative to controls. **G-H**) Summarized graphs showing reduced transition entropy (At 8 weeks, control = 5.32±0.03, n = 11, mitoPark = 5.28±0.05, n = 11, P = 0.56; At 14 weeks control = 5.22±0.06, n = 8, mitoPark = 5.23±0.06, n=8, P = 0.88; At 24 weeks control = 5.45±0.03, n = 7, mitoPark = 5.17±0.05, n = 9, P =0.0007; Mann-Whitney U test) and transition frequency (H, At 8 weeks control = 1.59±0.03, n = 11, mitoPark = 1.57±0.03, n = 11, P =0.4; At 14 weeks control = 1.56±0.04, n = 8, mitoPark = 1.48±0.05, n = 8, P = 0.38; At 24 weeks control = 1.76±0.06, n = 7, mitoPark = 0.85±0.05, n = 9, P = 0.0002; Mann-Whitney U test) of behavioral modules in mitoPark mice at 24 weeks, but not at earlier stages, relative to littermate controls.

In addition to the emergence of differently expressed behavioral modules over time, the difference in their usage was scaled up from the early to late stages of Parkinsonism. For example, the magnitude of reduction in the usage of fast syllables 1 and 3 of mitoPark mice relative to controls continued decreasing from 14 weeks to 24 weeks (see Table 2 for details). These observations are consistent with the concept that striatal neuronal activities modulate the vigor of movement that is closely linked to rapid DA release in the striatum ^7,23^.

### Time course of temporal organization of behavioral modules in progressive Parkinsonism

Next, we studied how progressive loss of DA affects the temporal organization of spontaneous locomotor activity. Plotting the expression of distinct syllables against time revealed that control mice exhibited dynamic and frequent transitions among behavior modules in locomotion (**Figure 3E**). However, such dynamic syllable transitions became less frequent in mitoPark mice, particularly at 24 weeks of age (**Figure 3E, G, H**). A few syllables, including syllable 0 (rearing), syllable 8 (grooming), and syllable 11 (slow walk) occupied long periods of time in mitoPark mice (**Figure 3E**). This conclusion was further supported by emerging differences in syllable transition probabilities between controls and mitoPark mice across ages (**Figure 3F**), manifesting as upregulated transition probability between a subset of syllables but downregulated transition probability between other syllables. The above data suggest that the naturalistic behavior of progressive mitoPark mice becomes more stereotypical and predictable as midbrain DA neurons progressively degenerate, recapitulating key findings from the 6-OHDA mice with complete DA depletion.

### L-DOPA effectively increases the velocity of behavioral modules in Parkinsonism but exerts a negligible effect on their usage

The mitoPark mice with advanced Parkinsonism are responsive to the L-DOPA administration ^9^. Next, we determined if L-DOPA can rescue the altered kinematics and usages of behavioral modules of mitoPark mice at 24 weeks. We intraperitoneally (i.p.) injected L-DOPA (10 mg/kg) and saline to mitoPark mice and age-matched controls, followed by monitoring their locomotor activities in an open field arena for 20 min. Analysis of conventional measurements of locomotor activity showed that L-DOPA administration effectively alleviated parkinsonian hypokinetic deficits of mitoPark mice (speed of locomotion, saline-injected control = 10.3±1.2 cm/sec, n = 9 mice; saline-injected mitoPark = 4.0±0.36 cm/sec, n = 9 mice, p = 0.004 vs saline-injected controls; mitoPark with L-DOPA = 14.4.9±2.97 cm/sec, n = 9 mice, P = 0.003 vs saline-injected mitoPark mice).

MoSeq identified various behavioral modules showing altered velocity and usage among groups. To determine the beneficial effects of L-DOPA, we identified a subset of behavior modules showing significant changes in their velocity and usage between saline-injected controls and mitoPark mice. Specifically, a group of fast behavioral modules, including run– and turn-like behaviors, showed decreased velocities between saline-injected mitoPark and saline-injected controls (**Figure 4A**). The velocities of most of these behavioral modules were significantly increased by L-DOPA administration. For example, the velocities of syllable 1 were 10.3±1.37 cm/sec in saline-injected controls (n= 5), 4.9±0.27 cm/sec in saline-injected mitoPark mice (n = 9), and 9.44±1.29 cm/sec in mitoPark with L-DOPA (n = 9) (saline-injected controls versus saline-injected mitoPark, P < 0.001; saline-injected mitoPark versus mitoPark with L-DOPA, P = 0.009, **Figure 4B**, see Table 3 for details).

**Figure 4.**
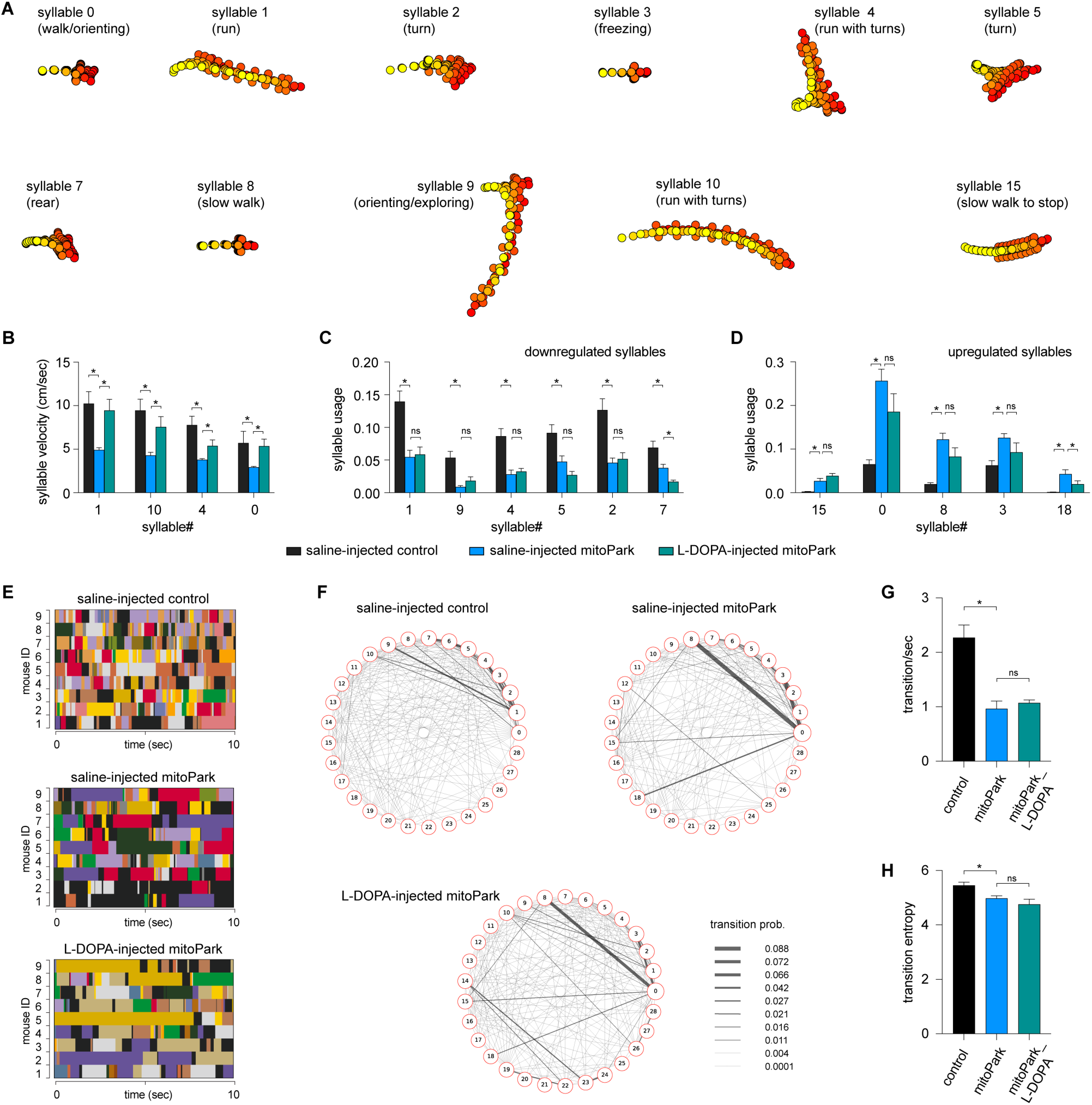
L-DOPA partially restores behavioral modules of mitoPark mice. **A**) Representative syllables showing significant changes in either velocities or usage between mitoPark mice and littermate controls. This subset of syllables was selected to study L-DOPA’s treatment efficacy. **B**) Summarized graph showing L-DOPA rescued the reduced velocity of behavioral syllables in mitoPark mice. **C-D**) Summarized graphs showing that L-DOPA affects the usage of neither the downregulated nor the upregulated behavioral syllables in mitoPark mice relative to the control. **E**) Representative plots showing the temporal organization of spontaneous behavior in controls (upper), mitoPark mice (middle), and mitoPark mice treated with L-DOPA (bottom) at 24 weeks. Different colors represent distinct behavior syllables. **F**) Transition graphs representing the changes in behavioral structure as an increase (red) or decrease (blue) in the probability of transition between numbered syllables. **G**) Summarized graphs showing reduced transition frequency (G, control = 2.27±0.23, n = 9; mitoPark = 0.96±0.14, n = 9, P = 0.0002 vs. control; mitoPark + L-DOPA = 1.07±0.05, n = 9, P > 0.05 vs. mitoPark) and transition frequency (H, control = 5.42±0.12, n = 9; mitoPark = 4.98±0.096, n = 9, P = 0.05 vs. control; mitoPark + L-DOPA = 4.76±0.19, n = 9, P > 0.05 vs. mitoPark) of behavioral modules in mitoPark mice at 24 weeks, which was not rescued by L-DOPA treatment.

Moreover, mitoPark mice exhibited both down– and up-regulated usage of behavioral modules. Surprisingly, L-DOPA treatment failed to restore the usage of downregulated modules in mitoPark mice (**Figure 4C**). When it comes to the upregulated behavior modules, the L-DOPA did not restore their frequency of expression either (**Figure 4D**). The above data suggest that L-DOPA administration increased the velocities of behavioral modules during spontaneous locomotory activity but had little effect on their usage in the Parkinsonian state.

Given the negligible effect of L-DOPA on syllable usage of the advanced mitoPark mice, we predicted that L-DOPA does not affect the transition probabilities and temporal organization of behavior modules in the Parkinsonian state. As predicted, we found that L-DOPA administration did not rescue the reduced behavior module transitions in mitoPark mice (**Figure 4E-H**). Specifically, neither the reduced transition frequencies (**Figure 4G**) nor the entropy (**Figure 4H**) in mitoPark mice was improved by L-DOPA administration. The above data suggest that the disrupted syllable usage and their temporal organization in Parkinsonian mice were not responsive to dopaminergic medication.

Altogether, the above results suggest that acute L-DOPA significantly improves the motor function of Parkinsonian mice and that its beneficial effect was mediated mainly by increasing the velocities of behavioral modules. L-DOPA administration produced little effect on the usage of behavior modules and their transitions during naturalistic locomotor behavior.

## Discussion

In the present study, we employed an unbiased and unsupervised approach to quantify the naturalistic locomotor behavior under Parkinsonian state. Using 6-OHDA mice with complete DA depletion, we demonstrate that complete loss of DA decreases the syllable velocities, disrupts the pattern of syllable usage, and reduces the transition probabilities among behavior modules (**Figure 1**). Correlation analysis further showed a stronger effect of DA depletion on the velocity and usage of behavioral modules with higher velocities (**Figure 1**). These observations are further confirmed and validated by using the mitoPark mice, a genetic and progressive model of Parkinsonism (**Figures 2 and 3**). The beneficial effects of L-DOPA on motor symptoms in PD have been well-established for decades. Therefore, we further analyzed the effects of L-DOPA on behavioral modules at the sub-second time scale. We demonstrate that L-DOPA effectively restored the decreased velocity but had a negligible effect on the disrupted usage of behavioral syllables of mitoPark mice (**Figure 4**). Together, our analyses support that hypokinetic symptoms in PD are associated with reduced movement velocity and abnormal usage of behavioral modules and that the therapeutic effects of L-DOPA are mediated by restoring velocities of syllables but not their temporal organization.

Spontaneous movements have been closely linked with the slow changes in tonic DA concentrations in the striatum ^24^. Recent work demonstrates that SN DA neuronal activity and the phasic striatal DA dynamics regulate movement initiation and generations of motor sequences during spontaneous behavior ^7,25,26^. Our observations from Parkinsonian mice are largely consistent with these earlier reports. Specifically, the bradykinesia-like hypokinetic motor deficits of mitoPark mice at the early motor stage with moderate striatal DA depletion are mainly driven by the reduced velocities of faster behavioral modules. As the SN DA neurons degenerate, the mitoPark mice gradually show decreases and increases in the usage of faster and slower syllables, respectively, from the early motor stage at 14 weeks to the advanced motor stage at 24 weeks (**Figures 2 and 3**). The combined effects of DA loss to syllable velocity and usage lead to the manifestation of severely impaired locomotor activity in both models of Parkinsonism.

Striatal DA transmission is critical in regulating goal-directed and spontaneous behavior by modulating postsynaptic cellular function, synaptic transmission, and plasticity processes within the corticostriatal circuits. Thus, the decreased velocity in Parkinsonism may be mainly driven by the decreased tonic DA levels in the striatum and can be rescued by L-DOPA administration. Moreover, we found that loss of DA downregulates the usage of several fast syllables and upregulates the usage of several slow syllables in both 6-OHDA mice and late-stage mitoPark mice. The altered usage of behavioral modules is likely to involve changes in striatal DA dynamics (e.g., phasic DA release) or other maladaptive changes in the cortico-striatal circuits that can’t be rescued by acute L-DOPA administration.

Taking advantage of the progressive nature of mitoPark mice, we showed that behavior modules with higher velocities (e.g., run– and walk-like syllables) were more vulnerable at early stages than those involving whole-body coordination (e.g., turning and rearing). Moreover, the magnitude of changes in the velocity and usage of the most vulnerable syllables increased from the early to advanced stages of mitoPark mice. This might reflect the exaggerated damage of cortico-basal ganglia circuits as the nigrostriatal pathway progressively degenerates. These data suggest that distal limb control was probably affected earlier (e.g., at 14 weeks), leading to a decreased expression of faster behavior modules at the early stages of nigrostriatal degeneration. Subsequently, proximal limb control and whole-body coordination were impaired at a later time point (e.g., 24 weeks), resulting in frequent pauses and fewer whole-body turns during movements that might contribute to the development of freezing of gait-like behaviors under parkinsonian state. Of relevance, the onset of motor symptoms in patients often starts from the distal limbs (e.g., fingers and feet) ^27^, which might affect their performance in skilled motor function or generate fast action sequences.

The literature on Parkinsonian animals has documented that nigrostriatal DA circuits are required for generating stereotypical behavioral sequences ^28^. The earlier studies focused on species-specific behavioral sequences in rodents (e.g., grooming) because they are readily identified by an experienced observer from recorded videos. Machine vision-based technologies make it possible to expand this research to less visible behavioral sequences during natural behavior. Using two mouse models of nigrostriatal degeneration, we documented that degeneration of the nigrostriatal projections alters the usage of distinct behavioral modules and decreases their transition probability in Parkinsonian states. This is consistent with earlier studies documenting deficits in motion sequences and switches between motor programs in PD patients ^29^.

Together, the progressive loss of DA does not alter behavioral modules in a unidirectional, uniform, and all-at-once manner. This observation further indicates the involvement of various biological mechanisms in regulating syllable velocity and usage and the therapeutic efficacy of L-DOPA under Parkinsonian state, including the roles of tonic versus phasic dopamine transmission by subtypes of nigral DA neurons ^25^, and potential contribution of the motor cortical network ^30^.

PD is featured by progressive loss of nigrostriatal DA projections, starting from the dopaminergic axons, followed by the loss of cell bodies ^31^. This “dying back” pattern of neurodegeneration has been recapitulated in several animal models, including the mitoPark mice ^8,32^. Thus, the compromised striatal DA levels in PD are expected to underlie the decreased velocity and usage of fast behavioral modules in both mouse models of Parkinsonism in the present study. L-DOPA administration can effectively restore tonic DA levels in the striatum, which results in the rescued velocity of behavioral modules in Parkinsonian mice. This is consistent with the observation that phasic DA transmission in the striatum does not modulate movement kinematics ^7,25^. Optogenetic stimulation of striatal dopaminergic axon terminals induces rapid phasic transmission and increases the acceleration frequency, triggering the initiation of locomotion. These observations indicate that phasic DA likely mediates the usages of behavioral modules with fast velocities. Given that L-DOPA administration can’t restore phasic DA transmission, it explains the failure of L-DOPA in rescuing the usage of behavioral modules in the late-stage mitoPark mice.

We demonstrated that L-DOPA rescued the decreased velocities of various behavior syllables of Parkinsonian mitoPark mice but had little effect on the usage and transitions of behavioral modules. Similar observations were reported by earlier studies in humans, showing that levodopa improved the slowed movement but did not alleviate the sequence effects in PD patients ^33^. Acute L-DOPA may restore tonic dopamine levels in the striatum, which play a key role in motor vigor and reinforcement learning ^34^. L-DOPA-induced increases in the tonic striatal DA levels likely underlie the restoration of motor velocity. Fast dopamine dynamics in the striatum works as reinforcement to promote the use and organization of behavioral modules ^7^. Thus, the disrupted behavioral sequencing in Parkinsonian mice is likely due to the impaired phasic DA release in the striatum that systematic L-DOPA injections can’t restore. However, we can’t exclude the involvement of other mechanisms, including maladaptive changes within and outside the striatum, in the abnormal usage of behavioral modules and sequencing ^35^.

In conclusion, the present work reveals key features of behavioral module changes in Parkinsonism. We demonstrated that loss of DA induces decreased velocity, abnormal usage, and fewer transitions of behavioral modules during spontaneous behavior, underlying the manifestation of hypokinetic deficits in PD. Further, we documented that acute L-DOPA-induced therapeutic effects are largely mediated by its effects on the velocity of behavioral modules but not their usages and sequences during spontaneous behavior. Further studies are warranted to better understand the biological mechanisms underlying impaired behavioral sequencing after loss of DA and design effective treatment to restore proper organization of naturalist behavior in the Parkinsonian state.

## Acknowledgement

Behavior recordings were conducted at the Van Andel Research Institute. Data analysis and manuscript preparation were performed and completed at Georgetown University Medical Center. This work was partially supported by research grants from the National Institutes of Health (R01NS121371, R21NS135545, H.Y.C; and R01NS133338, S.M.D. and H.Y.C.), Department of Defense Congressionally Directed Medical Research Programs (W81XWH-21-1-0943, H.Y.C.), and Parkinson’s Foundation (PF-IMP-1045313, H.Y.C.). This work was also funded in whole or in part by Aligning Science Across Parkinson’s (ASAP-020572) through the Michael J. Fox Foundation for Parkinson’s Research (MJFF). For the purpose of open access, the authors have applied a CC BY public copyright license to all Author Accepted Manuscripts arising from this submission.

## Author contributions

Conceptualization (DB, HYC); Investigation (DB, GS, HDC, LC); Formal analysis (DB, HYC); Validation (DB, LC, HYC); Writing – original draft (DB, HYC); Writing – Review & Editing (HYC); Supervision, Project administration and Funding acquisition (HYC).

## Availability of data and materials

The raw and analyzed during the current study have been deposited and are publicly available from Github (https://github.com/ChuLab-GU/behavior-analysis) and Zenodo (https://zenodo.org/records/14555868).

## Competing interests

The authors declare that they have no competing interests.

## Notes

https://zenodo.org/records/14555869

https://github.com/ChuLab-GU/behavior-analysis

### Competing Interest Statement

The authors have declared no competing interest.

### Summary of Updates

We revised the manuscript by adding more results and discussion. We also updated Key resources table to promote transparency and reproducibility.

